# *Dpp6* Knockout Mice Exhibit Increased Ethanol Conditioned Place Preference and Acute Ethanol-Induced Anxiolytic Behavior

**DOI:** 10.1101/2025.08.15.670589

**Authors:** Maribel Hernández, Amanda M Barkley-Levenson

## Abstract

The gene *DPP6* has been associated with behavioral phenotypes of alcohol use disorder (AUD) in recent human genome wide association studies. *DPP6* encodes an auxiliary subunit that modulates A-type voltage-gated potassium channels, particularly Kv4.2. To further assess the role of this gene in ethanol-related traits, we tested *Dpp6* knockout (KO) mice for ethanol conditioned place preference (CPP), locomotor activity, and ethanol-induced anxiolysis. Male homozygous KO mice (HOM) showed greater preference for the ethanol-paired context compared to wild type littermates (WT) and heterozygous KO mice (HET), while female mice showed no genotypic difference. HOM of both sexes exhibited greater novelty-induced hyperactivity in the CPP apparatus than HET and WT mice in the first two minutes. In a separate experiment, HOM mice showed enhanced locomotor activity following a 1.5 g/kg ethanol injection; however, they also displayed greater locomotor activity during habituation, suggesting basal locomotor differences. Following 1.5 and 2 g/kg injections, HOM mice exhibited EtOH-induced anxiolysis in the first 5 minutes, while the HET and WT mice did not. Lastly, HOM mice displayed a significant sedative response compared to WT animals following a 2 g/kg injection of ethanol. Ultimately, these findings validate a role for *Dpp6* in modulating ethanol’s rewarding, anxiolytic, and sedative effects in a sex-dependent manner.

## Introduction

As of 2023, 10.2% of the United States population aged 12 and older met criteria for Alcohol Use Disorder (AUD) within the past year (“2023 NSDUH Detailed Tables | CBHSQ Data,” n.d.). AUD is a complex disorder with both genetic and environmental factors contributing to its etiology. In fact, AUD is estimated to be roughly 50% heritable (Verhulst et al., 2015), and emerging evidence from large human genome-wide association studies (GWAS) have identified several novel genes associated with AUD phenotypes. More recently, the *dipeptidyl peptidase-like protein 6 (DPP6)* gene has been associated with measures of alcohol consumption (e.g., number of drinks per week), problematic alcohol use, and alcohol use disorder (AUD) measurement (Deak et al., 2022; Hade et al., 2021; Zhou et al., 2020), suggesting it may play a role in modulating alcohol-related behaviors.

*DPP6* encodes an auxiliary subunit that modulates the biophysical properties and surface expression of A-type voltage-gated potassium channels, particularly Kv4.2. Notably, *Dpp6* is most densely expressed in the hippocampus (HPC) and prefrontal cortex (PFC), regions that are both implicated in reward processing and learning (Chau et al., 2018; Iyer et al., 2025; Lasseter et al., 2010; LeGates et al., 2018a, 2018b; Sigurdsson and Duvarci, 2016). Behavioral phenotyping of *Dpp6* knockout (KO) mice has revealed impairments in tasks related to recognition and spatial learning and memory, as demonstrated in paradigms such as the novel object recognition and Morris water maze tasks (Lin et al., 2018). Additionally, significant changes in locomotor behavior and weight have been reported in mice with a *Dpp6* deletion. More specifically, KO mice tend to show increased locomotor activity and exhibit lower body and brain weights than wild types (WT) (Lin et al., 2022, 2018). However, no preclinical studies have assessed the role of *Dpp6* in any alcohol-related phenotypes.

Ethanol (EtOH) conditioned place preference (CPP) is a well-established associative learning paradigm, whereby EtOH is paired with a specific context to elicit an unconditioned stimulus response. In this case, we can assess the rewarding/aversive properties of EtOH, which contribute to the development of AUD. Several brain regions have been implicated in CPP acquisition and expression, such as the bed nucleus of the stria terminalis (BNST), nucleus accumbens (NAc), PFC, and HPC (Dong et al., 2006; Hitchcock and Lattal, 2018; Pati et al., 2019a; Zavala et al., 2003). *Dpp6* KO mice show changes in dendritic morphology and excitatory neurotransmission in the HPC (Lin et al., 2013), which we hypothesized could result in changes in EtOH CPP expression.

Here, we sought to validate the role of *Dpp6* in modulating sensitivity to ethanol’s rewarding and locomotor effects through two complementary experiments. Experiment 1 used an EtOH CPP protocol to determine whether *Dpp6* KO mice exhibit enhanced conditioned preference for an EtOH-paired context compared to WT controls. Experiment 2 assessed whether *Dpp6* KO alters the acute locomotor and anxiolytic response to a stimulatory (1.5 g/kg) or mildly sedative (2 g/kg) dose of EtOH in each of the genotypes. Together, these experiments highlight that the global KO of *Dpp6* alters behavioral responses to EtOH.

## Materials and Methods

### Animals

A total of 66 adult *Dpp6* homozygous KO (HOM), heterozygous KO (HET), and wild type (WT) littermate mice were used in Experiment 1 (both sexes; postnatal day [PND] 57–152; n=4-17/sex/genotype). Experiment 2 used 20 adult mice of all three genotypes (both sexes; PND 74–101; n=4-8/sex/genotype). All animals came from an in-house breeding colony established from B6.Cg-Dpp6*^tm1.1Dahn^*/J cryorecovered breeders (Jax stock # 017972; Su et al. 2011) and maintained with HET x HET breeder pairs. All mice were genotyped via TransnetYX (Memphis, TN) using validated protocols. Mice were group-housed (2–5 per cage) in large Plexiglas cages equipped with a paper house and plastic upper-level compartment for enrichment and maintained on a 12:12 h light/dark cycle (lights on at 07:00). Food (PicoLab 5L0D) and water were available *ad libitum*. All procedures complied with institutional and national ethical standards and were approved by the University of New Mexico Health Sciences Center Institutional Animal Care and Use Committee.

### Place conditioning and open field apparatus

CPP testing was conducted using a two-chambered apparatus (Med Associates, Inc., VT) fitted into a clear Plexiglas chamber (27.3 × 27.3 × 20.3 cm). The CPP insert consisted of two tactilely distinct compartments (12.7 cm x 25.4 cm) separated by a black wall with a removable guillotine door. One side featured a stainless-steel bar floor (approximately 5 mm between the center of each bar), and the other a stainless-steel grid mesh floor; both were flanked by clear walls. The open field test used the same Plexiglas chamber without the CPP insert. Locomotor and positional data were recorded using the Med Associates Activity Monitor system. Chambers were cleaned with 10% isopropyl alcohol between animals to eliminate olfactory cues.

### Drugs and solutions

0.9% sterile saline (UFC Bio, Amherst, NY, via Amazon) and 20% EtOH (v/v in saline) were administered intraperitoneally (i.p.) immediately prior to placement in the CPP chamber. Conditioning sessions alternated between 10 ml/kg saline or 2 g/kg EtOH injections. During the open field experiment, mice received a 1.5 g/kg and 2 g/kg EtOH (in 20% v/v in saline) injection, and the saline volume equivalent of a 1.5 g/kg injection. All solutions were prepared daily prior to behavioral testing.

### Experiment 1: Ethanol conditioned place preference

The CPP procedure consisted of a pretest, conditioning trials, and preference tests. On the pretest day, mice were moved into the testing room, weighed, and allowed to acclimate for at least 60 min. They were then injected with saline and immediately placed in the center of the apparatus and allowed to explore freely for 30 min to assess initial chamber preference. Due to an average group baseline preference of 60.17% ± .012 for the grid floor chamber, a biased conditioning approach was used where each animal was assigned to receive EtOH in their initially non-preferred chamber (>50%) as described in Cunningham, Ferree, and Howard (2003). Any animal showing an initial chamber preference ≥80% would be excluded from testing, though no mice in this experiment reached this initial preference cutoff. Five days after the pretest, conditioning began. During conditioning trials, the guillotine door was inserted to limit access to a single chamber, and animals were injected with the assigned drug and then immediately placed in the corresponding chamber for five minutes. For example, on day 1, animals were all placed in the bar floor chamber following either 2 g/kg or 10 ml/kg saline injection depending on whether the bar floor type was their conditioned stimulus (CS+) or unconditioned stimulus (CS-), respectively. The following day, all animals were placed in the chamber with the grid floor type and received injections according to their conditioning group. Five-minute conditioning trials were used as this is a standard protocol for ethanol CPP, and previous studies have determined that this trial duration produces a stronger preference compared to longer trials (e.g., Cunningham and Prather, 1992). Four days of conditioning trials were given each week for three weeks, alternating between saline and ethanol trials, with one trial given per day. On the preference test days (testing days 5, 12, and 19), all animals were injected with saline and placed in the center of the testing apparatus and allowed to freely explore both chambers to assess floor preference (30-minute test). The entire procedure was conducted 5 days per week with a two-day break between each conditioning and test cycle (Cunningham et al., 2006). Time spent in each chamber and distance traveled were automatically recorded on each test day. We computed a CPP change score for each preference test (% time in EtOH chamber on test day - % time in EtOH chamber on pretest).

### Experiment 2: Open field activity

Locomotor activity was recorded on three consecutive days in 30-minute sessions. On days 1 and 2, mice received a saline injection prior to placement in the open field chamber. On day 3, mice were injected with 1.5 g/kg ethanol (20% v/v in saline). A follow-up test was conducted one week later, following a 2 g/kg ethanol injection, during which activity was recorded over 60 minutes. Distance traveled (cm) and time spent in the center zone (14.45 cm x 14.45 cm) were automatically recorded.

### Data Analysis

All data were analyzed using SPSS software (SPSS, Version 30, Chicago, IL) and graphed and analyzed with GraphPad Prism software (GraphPad Prism, v. 10.5.0, La Jolla, CA). Animals were excluded from analyses on a given day if the Med Associates program did not track their behavior correctly. No animals were removed from the entire study as these issues were resolved for testing the following day, and mixed effects analyses were used. Repeated measure two-way or three-way analysis of variance (RM ANOVA) tests were used to assess test*genotype*sex or time*genotype*sex interactions in cases where there were no missing data points. If there was no main effect of sex or significant sex interactions, data were collapsed. Significance values were set at p < 0.05. All data were assessed for normality using the Shapiro-Wilks Test, and if normality was violated, non-parametric tests were used. For repeated-measures data, violations of Mauchly’s test for sphericity were corrected using Greenhouse–Geisser estimates.

## Results

### Experiment 1: EtOH Conditioned Place Preference

#### Only HOM males show significant increase in time spent in the EtOH-paired chamber across preference tests

This experiment tested whether the global homozygous or heterozygous knockout of *Dpp6* would alter EtOH CPP. Figure 1 shows CPP change scores (% time in CS+ on test N – % time in CS+ on pretest) using three-way RM ANOVA of test, sex, and genotype. We found a significant main effect of test, [F(1.73, 103.91) = 11.2, p < .001], and significant interactions of test*sex, [F(2, 120) = 7.7, p = .001], and test*sex*genotype, [F(4, 120) = 5.47, p < .001] (Figure 1A-B). To probe the three-way interaction, we ran separate two-way RM ANOVAs of test and genotype for each sex. For females, there were no significant main effects or interactions (Figure 1A). In males, a two-way RM ANOVA revealed a significant main effect of test, [F(1.58, 56.83) = 19.44, p < .001], and a significant interaction of test*genotype, [F(3.16, 56.83) = 5.79, p = .001] (Figure 1B). Tukey’s multiple comparisons tests confirmed significant increases in CPP score in HOM males from test 1 to test 2 (p = .02) and from test 1 to test 3 (p = .012), as well as between WT and HOM males on tests 2 (p = .016) and 3 (p = .004, HOM > WT). These data suggest that CPP change score was significantly increased in HOM males only, consistent with the raw CS+ preference scores.

**Figure 1.**
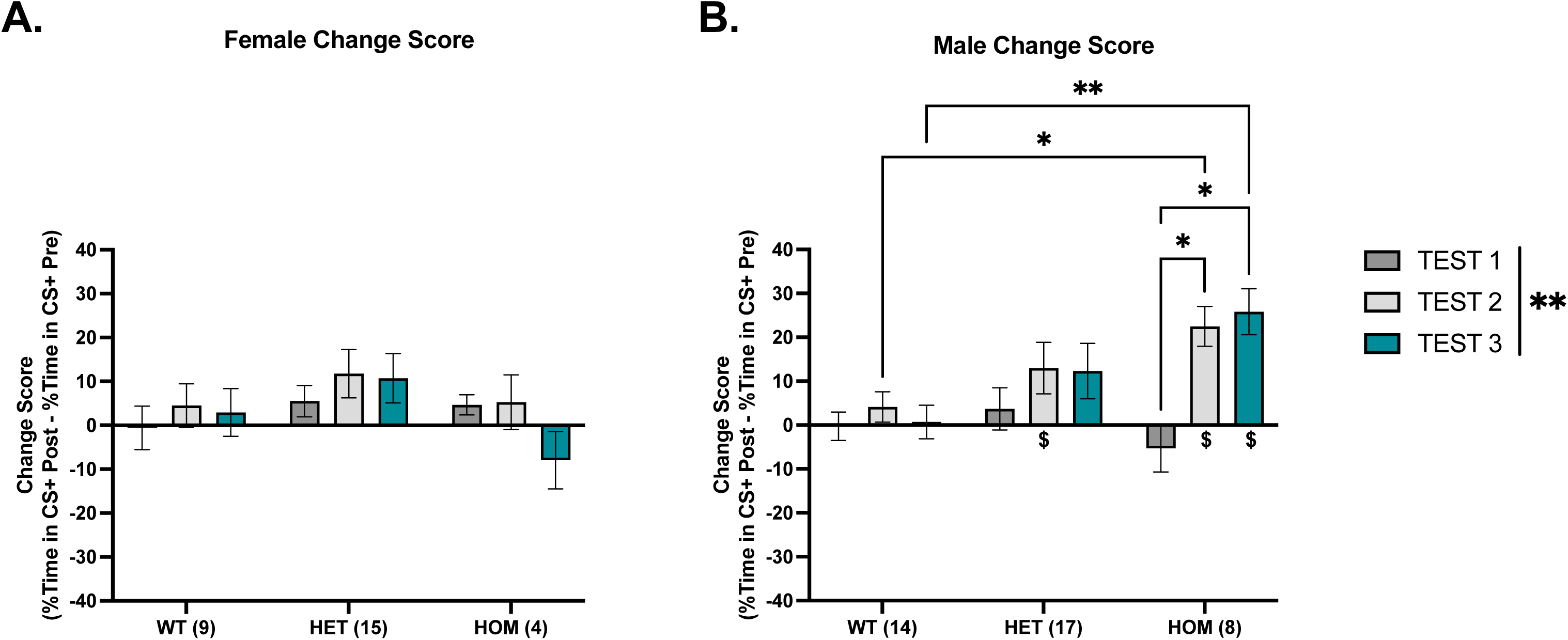
CS+ change scores for *DPP6* WT, HET, and HOM mice. A) Females did not show changes in their change score or in their ability to express CPP. B) Male KOs show significantly increased CPP change scores after conditioning on test 2 and test 3, both of which are significantly higher as compared to WTs. When assessing their CPP expression, only HET males on test 2 and HOM males on test 2 and 3 showed significant change from zero. WT Female (n = 8); HET Female (n=15); HOM Female (n= 4); WT Male (n= 14); HET Male (n= 17); HOM Male (n= 8) (*p < .05) (**p < .01) (^$^p < .05, change from zero) (^$$^p < .01, change from zero)

#### Only HET and HOM Males Show Significant EtOH CPP Expression

One-sample t tests of CPP change scores (vs. zero) were used to assess the development of significant EtOH CPP expression. In females (Figure 1A), no group showed significant effects, though HETs showed trends at test 2 (p = .051) and test 3 (p = .077). In the males (Figure 1B), HETs showed significance at test 2 (M = 12.98%, SD = 24.199%), [t(16) = 2.2, p = .042], and a trend at test 3 (p = .068). HOM males showed significant effects at test 2 (M = 22.47%, SD = 12.87%), [t(7) = 4.94, p = .002], and test 3 (M = 25.84%, SD = 14.85%), [t(7) = 4.92, p = .002]. Taken together, we can confirm that the HOM males and the HET males showed significant changes in preference for the CS+ context after two and three rounds of conditioning.

### Experiment 2: Open Field Activity

#### HOM Mice Display Novelty-Induced Hyperactivity

Next, we assessed locomotor activity during the pretest (Figure 2A). A three-way RM ANOVA of time (30 minutes), sex, and genotype revealed significant main effects of time, [F(1.14, 68.40) = 931.95, p < .001], and sex, [F(1, 60) = 10.09, p = .002, F > M], and a significant interaction of time*sex, [F(29, 1740) = 9.5, p < .001]. When collapsed by genotype, a two-way RM ANOVA of time and sex revealed a significant main effect of time, [F(16.66, 1067) = 58.20, p < .001], and sex, [F(1, 64) = 11.97, p < .001, F > M], and no significant interactions.

**Figure 2.**
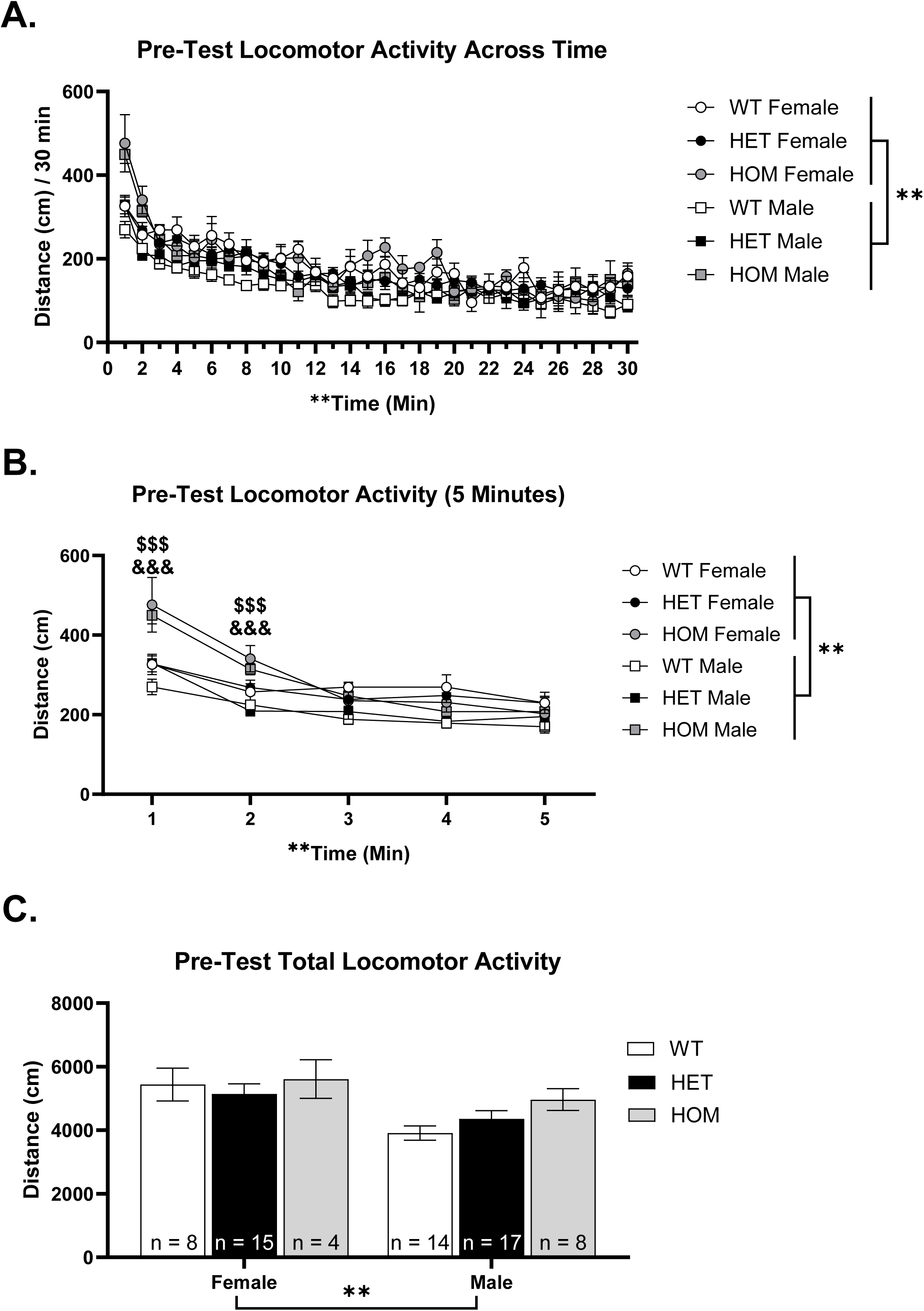
Pretest locomotor activity in Dpp6 WT, HET, and HOM females and males. A) pretest locomotor activity changed across the 30-minute session time and females moved around more than males. B) In the first 5 minutes of the session, both female and male HOM KOs exhibited greater locomotor activity in the first two minutes compared to WT and HET animals. Females also moved more than males in general. C) Total locomotor activity only showed a significant difference between females and males, suggesting that the genotypes move similarly. WT Female (n = 8); HET Female (n=15); HOM Female (n= 4); WT Male (n= 14); HET Male (n= 17); HOM Male (n= 8) (*p < .05) (**p < .01) (^##^p < .01, ME sex) (^$$$^p < .001, HOM vs WT) (^&&&^p < .001, HOM vs HET)

To assess potential genotypic differences in novelty-induced hyperlocomotion, we analyzed the first five minutes of the test on Day 1. This early time point was selected because the initial 5–10 minutes of open field exposure are widely considered the most sensitive period for capturing novelty-driven exploratory behavior (Gould, 2009). This focused analysis of the first five minutes revealed significant main effects of time, [F(4, 240) = 20.13, p < .001], and sex, [F(1, 60) = 10.03, p = .002], and a significant interaction of time*genotype, [F(8, 240) = 4.01, p < .001] (Figure 2B). There was a trend towards a time*sex*genotype interaction (p = .06). To follow up on the significant time*genotype interaction, we collapsed across sex. A two-way RM ANOVA, confirmed significant main effects of time [F(4, 252) = 122.8, p < .001] and genotype, [F(2, 63) = 5.264, p = .001], and a time*genotype interaction, [F(8, 252) = 11.35, p < .001]. Tukey’s multiple comparisons tests indicated the HOMs were significantly more active than WTs and HETs at minute 1 (both p’s < .001), and minute 2 (HOM > WT, p = .001; HOM > HET p < .001). However, total locomotor activity (30 minutes) showed only a main effect of sex, [F(1, 60) = 10.05, p = .002, F > M], suggesting that the genotypic difference in locomotor activity is due to the initial novelty of the context (Figure 2C).

#### HOMs Show Greater Locomotor Activity on Saline Injection Days in Open Field Test

Figure 3 shows locomotor activity on each day of the open field test. On habituation day 1, a three-way RM ANOA of time (30 minutes), sex, and genotype revealed main effects of time, [F(9.99, 289.63) = 30.06, p < .001], sex, [F(1, 29) = 8.14, p = .01, F > M], and genotype, [F(2, 29) = 13.68, p < .001, HOM > HET and WT], and significant interactions of time*sex, [F(9.99, 289.63) = 2.46], and time*genotype, [F(19.97, 289.63) = 2.08, p = .01] (Figure 3B). To probe the interactions, we collapsed on either sex or genotype. A two-way RM ANOVA of time and sex revealed a significant main effect of time, [F(8.5, 280.6) = 29.69, p < .001], sex, [F(1, 33) = 4.25, p = .047, F > M] and a time*sex interaction, [F(8.50, 280.6), p = .017]. Tukey’s multiple comparisons tests indicated that females were more active than males at minute 1, 3, 5, 13, 23, 24, and 30 (p’s ≤ .049). Next, a two-way RM ANOVA of time and genotype revealed a significant main effect of time, [F(9.33, 298.47) = 28.94, p < .001], genotype, [F(2, 32) = 9.84, p < .001, HOM > HET and WT], and an interaction of time*genotype, [F(18.66, 298.47) = 1.86, p = .017]. Tukey’s multiple comparisons tests revealed elevated activity in HOMs vs. WTs at multiple time points: 1-3, 10, 11, 14, 18, 19, and 22 (p’s ≤ .04) and HETs at minute 28 (p = .02). Next, a two-way ANOVA of sex and genotype on total locomotor activity (30 minutes) revealed a main effect of sex, [F(1, 29) = 8.144, p = .008, F > M], and genotype, [F(2, 29) = 13.68, p < .001 HOM > HET and WT] (Figure 3C), suggesting that females show greater locomotor activity than males and that HOMs show greater locomotor activity in a 30-minute test in a novel context.

**Figure 3.**
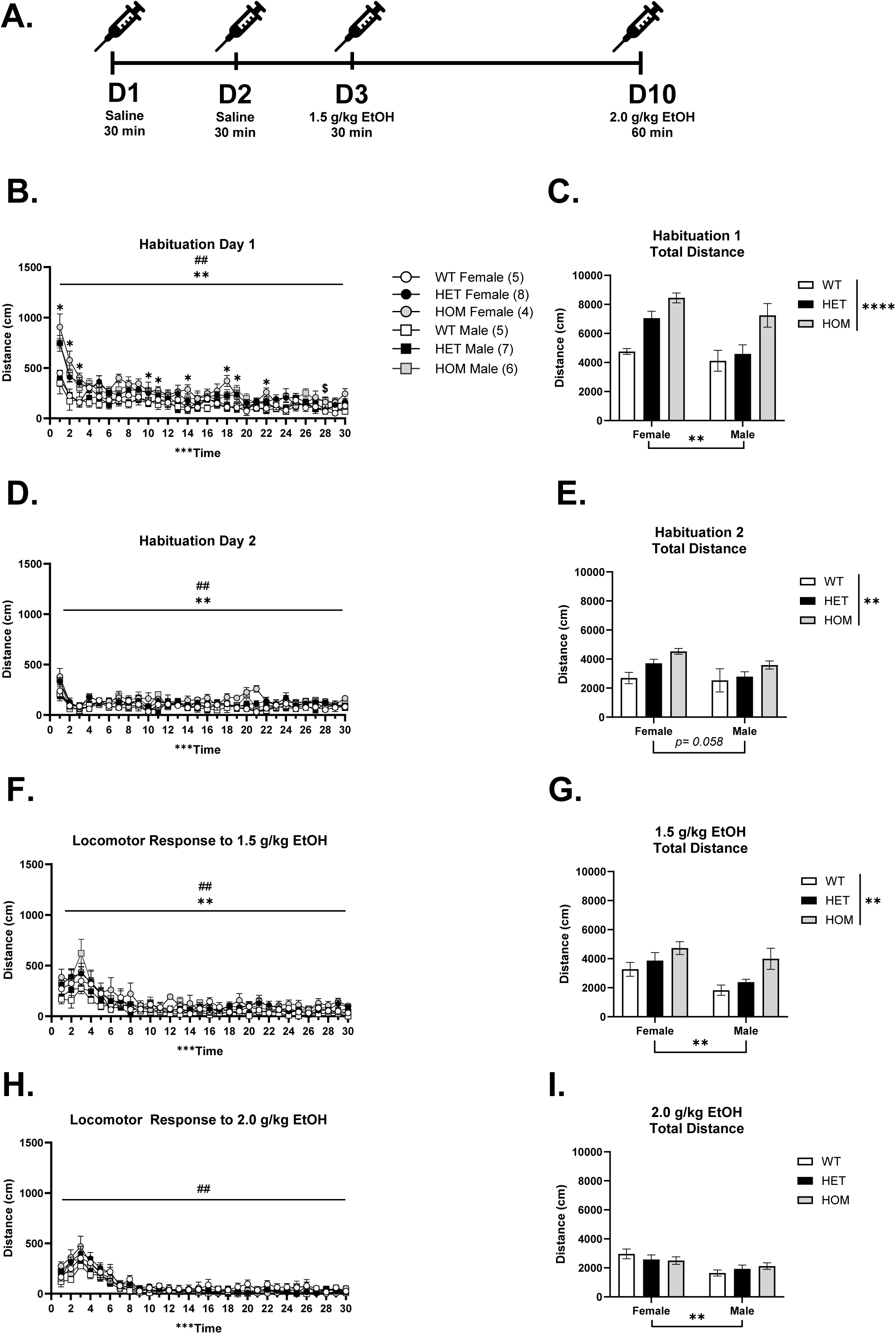
Assessment of 30-minute locomotor activity each day. A) A timeline of Experiment 2. B) Habituation day 1 showed significant effects of sex, genotype, and time. C) Total locomotor activity showed main effects of sex and genotype. D) Habituation day 2 also showed significant effects of sex, genotype, and time. E) Total locomotor activity on day 2 shows a significant effect of genotype. F) An acute injection of 1.5 g/kg EtOH showed significant effects of sex, genotype, and time. G) Total locomotor on day 3 showed significant effects of sex and genotype. Lastly, H) an injection of 2 g/kg on day 10 showed significant effects of sex and time but not genotype. Similarly, I) shows a main effect of sex only. WT Female (n = 5); HET Female (n=8); HOM Female (n= 4); WT Male (n= 5); HET Male (n= 7); HOM Male (n= 6); (***p < .001) (^##^p < .01, ME of sex) (**p < .01 and ME genotype across time)

On habituation day 2, one HET male was excluded from analysis due to equipment malfunction. A three-way RM ANOVA of time, sex, and genotype revealed significant main effects of time, [F(29, 812) = 31.42, p < .001], sex, [F(1, 28) = 7.39, p = .01, F > M], and genotype, [F(2, 28) = 5.35, p = .01, HOM > HET and WT], with a trend toward a time*genotype interaction (p = .06). These data suggest that on the second day of the saline injection, females moved more than males and, overall, there were genotype differences across time (Figure 3D). Next, a two-way ANOVA of sex and genotype on total locomotor activity showed a genotype effect, [F(2, 28) = 5.48, p = .01], and a trend toward a main effect of sex (p = .06; Figure 3E). A post hoc Tukey’s multiple comparisons test revealed that HOM mice had significantly greater total distance traveled than WT mice (p = .007).

#### HOMs Show Greater Locomotor Activity Compared to WTs Following a 1.5 g/kg EtOH Injection but not a 2 g/kg EtOH Injection

On the first EtOH treatment day, a three-way RM ANOVA of time, sex, and genotype revealed significant main effects of time, [F(6.69, 194.09) = 32.97, p < .001], sex, [F(1, 29) = 8.17, p = .01, F > M], and genotype, [F(2, 29) = 5.58, p = .01, HOM > WT]. These data suggest that an acute injection of 1.5 g/kg ethanol produces a significantly more stimulatory response in females compared to males and that there are significant differences between genotypes (Figure 3F). Next, a two-way ANOVA of total locomotor activity showed similar main effects of sex, [F(1, 29) = 8.17, p = .01, F > M], and genotype, [F(2, 29) = 5.58, p = .01, HOM > WT] (Figure 3G), with HOM mice having greater activity compared to WTs, and females compared to males.

We recorded locomotor activity for 60 minutes on Day 10 to capture potential behavioral differences (e.g. recovery from sedative effects) at a later time point following a higher dose of EtOH. However, we focused our analyses on the first 30 minutes to remain consistent with the other test days. Following a 2 g/kg injection of EtOH, a three-way RM ANOVA of time (first 30-minutes), sex, and genotype revealed significant main effects of time, [F(6.07, 176.06) = 50.95, p < .001], and sex, [F(1, 29) = 10.74, p = .003, F > M] (Figure 3H). In the first 30 minutes post-injection (Figure 3I), a two-way ANOVA of total locomotor activity revealed a main effect of sex, [F(1, 29) = 10.74, p = .003, F > M], and no other significant effects, suggesting that at a higher dose locomotor activity was similar across all genotypes. Locomotor activity during the full 60 minutes can be seen in Figure S3.

#### HOMs Show Sedative Response to 2 g/kg EtOH Injection

Change scores were calculated for each EtOH dose using day 2 (saline) as baseline and two-way ANOVAs of sex and genotype were run on the change scores at each dose. Data were collapsed on sex if no significant main effects or interactions were present. At the 1.5 g/kg EtOH dose, a one-way ANOVA did not yield any significant effects, suggesting that the 1.5 g/kg dose did not induce a stimulatory response in any genotype, and the greater locomotor activity at this dose in the HOMs compared to WT does not reflect a greater stimulatory response (Figure 5A). At the 2 g/kg EtOH dose, when collapsed by sex, a one-way ANOVA confirmed a genotype effect, [F(2, 31) = 4.702, p = .017]. Specifically, there was a significant difference between WTs and HOMs (p = .018), suggesting that HOMs do display a greater sedative response in the first 30 minutes compared to WTs (Figure 5B).

**Figure 4.**
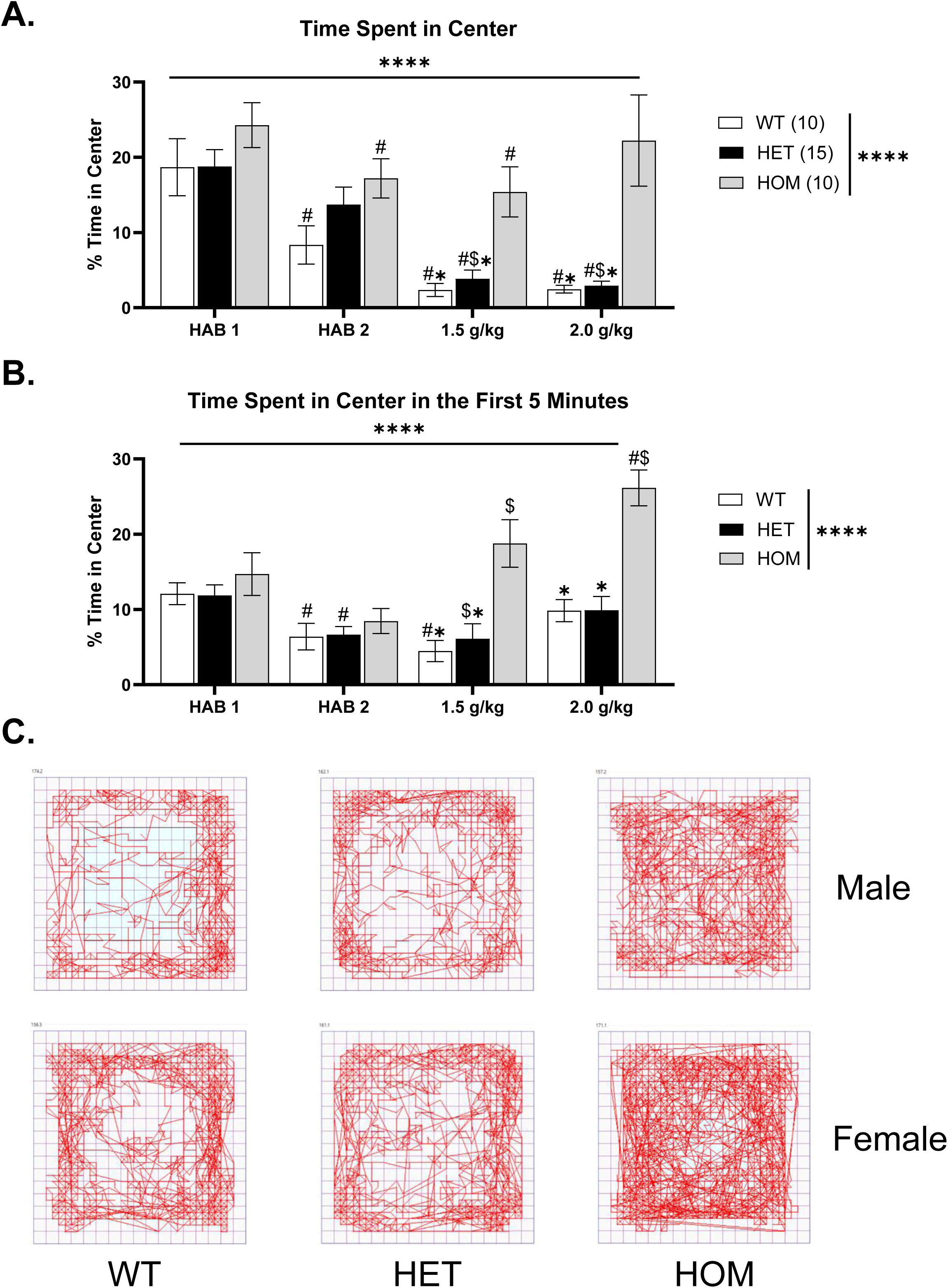
Anxiety-like behavior across test days. A) The percent time spent in the center was significantly difference between genotypes across days. HOMs, WTs, and HETs showed decreased anxiolytic behavior compared to HOMs on the EtOH injection days. B) When we looked at the first five minutes, we found that genotypes were different across testing days. Similarly, WTs and HETs were significantly different from HOMs on days when EtOH was administered. C) Representative images of the locomotor activity seen in each genotype on day 3 (1.5 g/kg injection; ^#^p < .05, different from Hab 1) (^$^p < .05, different from Hab 2) (*p < .05, different from HOMs in the same group).

**Figure 5.**
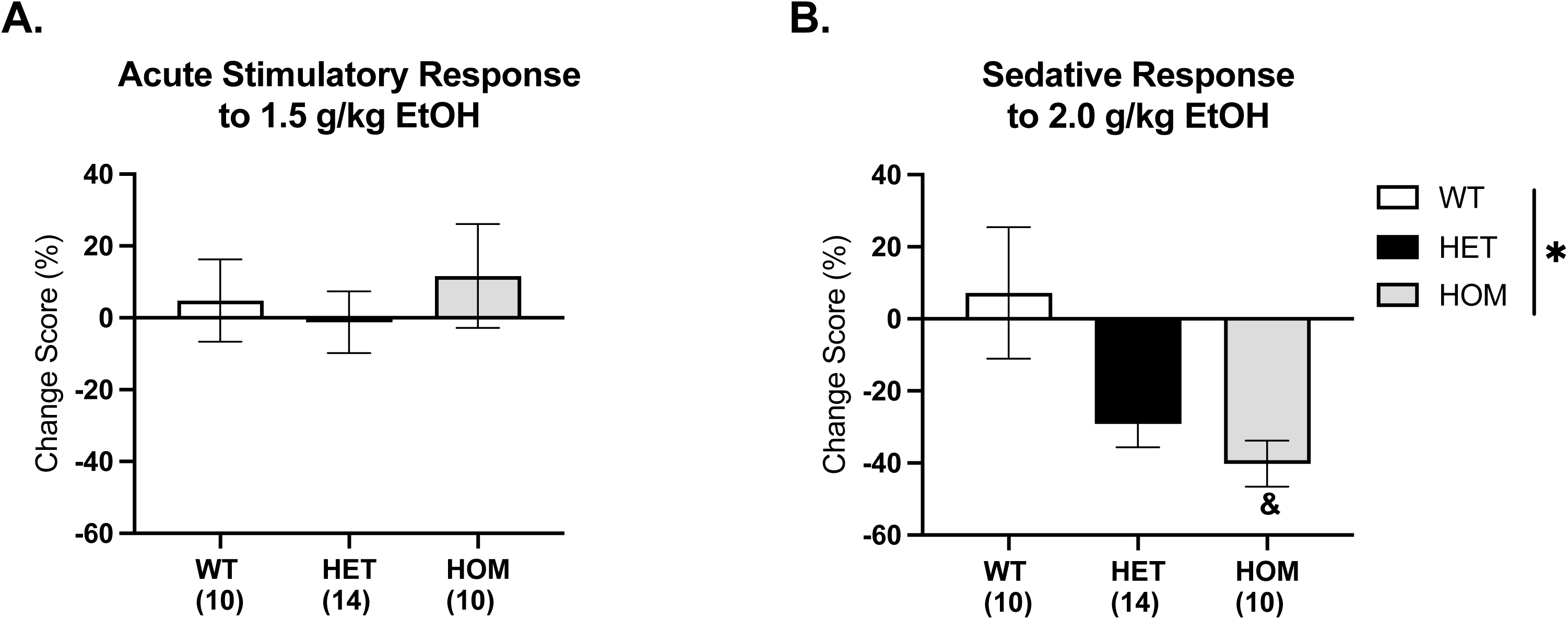
Change scores of EtOH injection days from habituation day 2. The change scores were calculated based on the first 30 minutes of each test. There were no main effects of sex, so groups were collapsed. A) 1.5 g/kg injection did not show any significant difference between sex or genotype. B) A 2 g/kg EtOH injection resulted in a main effect of genotype. Specifically, the HOMs showed a greater sedative response compared to WTs. (*p < .05) (^&^p < .05, different from WT)

#### Only HOMs Show EtOH-Induced Anxiolytic Behavior

Percent of time spent in the center of the Open Field Activity chambers across injection days can be seen in Figure 4. One HET male was again excluded from analysis on this day due to equipment malfunction. When collapsed by sex, a mixed-effects analysis revealed main effects of day, [F(1.917, 60.71) = 27.64, p < .001], genotype, [F(2, 32) = 8.868, p = .001, HOM > HET and WT], and a day*genotype interaction, [F(6, 95) = 3.942, p = .002] (Figure 4A). Tukey’s multiple comparisons test revealed HOMs spent significantly greater time in the center than WTs (p = .01) and HETs (p = .02) on day 3 (1.5 g/kg) and day 10 (2 g/kg; HOM > WT, p = .02; HOM > HET, p = .03), suggesting a genotypic-specific increase in anxiolytic response to ethanol. Within-genotype comparisons showed significant changes across days. In WTs, time spent in the center was lower on days 2, 3, and 10 compared to day 1 (p’s ≤ .01). In the HETs, time spent in the center was lower on days 3 and 10 compared to both day 1 and day 2(p’s ≤ .001). Lastly, HOMs spent significantly more time in the center on day 1 vs. day 2 (p = .001) and day 1 vs day 3 (p = .03). Thus, there were no differences between time spent in the center on EtOH injection days compared to the second saline injection day, which suggests that center time was not significantly different from the baseline day (Figure 4A) when looking at the data for the entire 30-minute session.

Because of the genotypic differences in locomotor activity during the initial minutes of testing, we decided to look at the change in percent time spent in the center in the first five minutes (Figure 4B). Further, this was done to avoid the potential confound of any sedative effect on locomotor activity (Bocarsly et al., 2019; Lister, 1987; Rose et al., 2013). When collapsed by sex (no significant effect or interactions), a mixed effects analysis of day and genotype revealed significant main effects of day, [F(2.398, 75.95) = 12.54, p < .001], and genotype, [F(2, 32) = 15.96, p < .001, HOM > HET and WT], and a significant interaction of day*genotype, [F(6, 95) = 5.561, p < .001]. Tukey’s multiple comparisons test indicated significant differences between WTs and HOMs at the 1.5 g/kg (p = .003) and 2 g/kg dose (p = .01). There were also significant differences between HETs and HOMs at the 1.5 g/kg (p = .01) and 2 g/kg dose (p < .001), with HOMs having greater center time than the other genotypes. There were also significant differences between days in each group. In WTs, there were differences between day 1 vs. day 2 and day 3 (p’s ≤ .04). In HETs, there were differences between day 1 vs. day 2 and 3 (p’s ≤ .04). Finally, in HOMs, there were differences between day 1 vs. day 10 (p = .01), day 2 vs. day 3 (p = .03) and day 2 vs. day 10 (p = .001), suggesting that in the first 5 minutes of the session only HOMs exhibited an EtOH-induced anxiolytic response at both doses tested. Representative images of locomotor activity in the chamber on day 3 (1.5 g/kg EtOH injection) can be seen in Figure 4C.

## Discussion

To our knowledge, this study is the first to validate the involvement of *Dpp6* in alcohol-related phenotypes and reveals several key differences between WT and KO mice. In Experiment 1, HOM males developed significant ethanol CPP, but the other genotypes and the HOM females did not. Additionally, HOM males and females exhibited significant increases in their initial locomotor activity during the pretest compared to WT mice, suggesting novelty-induced hyperactivity. In Experiment 2, we replicated this novelty-induced hyperactivity in the HOMs, and we saw greater total locomotor activity across days compared to WT mice. Interestingly, in HOM males and females, both a 1.5 g/kg and a 2 g/kg injection of EtOH significantly increased the percentage of time spent in the center of the open field during the first 5 minutes of the test test compared to the other genotypes. This suggests a greater anxiolytic response to EtOH in these mice in the first 5 minutes after injection, though this effect did not persist when analyzing the entire 30-minute session. When we compared the difference in locomotor activity between each EtOH injection day and baseline (habituation day 2), we found that the 2 g/kg dose produced a significant sedative response in the HOMs as compared to the WTs, whereas no genotype showed a significant stimulatory effect at the 1.5 g/kg dose. We also assessed weights in these mice because a loss of *Dpp6* has been previously associated with lower body weight (Lin et al., 2022, 2018). Specifically, Lin et al. (2018) found that *Dpp6* loss resulted in significantly lower body weights and brain weights at different developmental times. Here, we did not find significant differences between our WT and KO littermates (Figure S1 & S3); however, we did see a trend in the previously reported direction.

In Experiment 1, we found a significant CPP response on test 2 and 3 in the male HOMs, and on test 2 in the male HETs (Figure 1B), suggesting that the male HOMs are more sensitive to the rewarding properties of EtOH. A potential limitation of this study is that the WTs did not develop a preference for the EtOH-paired chamber, which could be explained by their C57BL/6J (B6) background. Several studies have highlighted that B6 mice are not as sensitive to the rewarding properties of EtOH as compared to other strains, and only modestly show EtOH CPP (e.g. Cunningham and Shields, 2018). However, the male HOMs were able to form a significant context-drug association despite the same B6 background. Interestingly, we found that female HOMs did not condition to the EtOH-paired chamber, which suggests that *Dpp6* might be involved in a sex-dependent manner in associative or context-dependent learning. Alternatively, this effect could also be due to general sex differences in sensitivity to the rewarding properties of alcohol; however, this remains unexplored and should be considered moving forward. It should be noted that we had a smaller number of female HOMs in this study compared to the other sex and genotype groups, which is a limitation. It is possible that we are therefore underpowered to detect the genotype by sex interaction, and the sex-specific findings might not persist with a larger sample size. Thus, an additional future experiment with larger samples sizes is needed in order to fully understand the sex difference seen here.

We observed that male HOMs increased their preference for the EtOH-paired chamber following the second conditioning block, suggesting that additional conditioning days were required to strengthen the context-drug association. Previous studies have demonstrated that test days can act as extinction trials because repeated exposure to the conditioned context without the drug can lead to attenuation of preference across tests (Yahyavi-Firouz-Abadi and See, 2009). However, we found an enhanced response in HOM males across preference tests, rather than evidence of extinction. This is likely due to the four days of conditioning trials between each test day, which presumably counteracted any extinction learning occurring on the preference tests.

We opted to use a biased conditioning approach in this study because we observed an initial group preference across all genotypes for the grid floor type (average pretest preference 60.17% grid vs 39.83% bar). Interpretation of the results from biased EtOH CPP paradigms must consider whether the CPP response is due to the rewarding effects of EtOH or due to EtOH’s ability to remove an aversive state in the non-preferred context (e.g. anxiolytic effect) (Cunningham et al., 2003; Schenk et al., 1985). However, Cunningham, Ferree, and Howard (2003) were able to find significant conditioned place preference in both an unbiased and biased protocol using the same EtOH dose, which suggests that one cannot conclude that this effect is only due to EtOH’s anxiolytic effects, but rather a combination of both the rewarding and anxiety-reducing properties of EtOH. Regardless, if the CPP expression seen in our biased approach did solely rely on the initially non-preferred side becoming “less aversive” or “less anxiety-inducing” we can speculate that, in HOM males, there may be alterations in anxiety-related circuitry. However, this is beyond the scope of the present studies and should be considered as a future direction.

Another consideration in the interpretation of these findings is the relationship between EtOH-induced locomotor activation and CPP expression. We did not see evidence of EtOH-induced locomotor stimulation in either experiment, and in fact found a significant sedative response in the HOMs at the 2 g/kg EtOH dose in the open field experiment, which is the same dose used for EtOH CPP in Experiment 1. Importantly, research suggests that EtOH locomotor stimulation and CPP are dissociable and that EtOH CPP can be present without locomotor stimulation and vice versa (Acevedo et al., 2013). High levels of locomotor activity during the preference test have been shown to obscure CPP expression (Gremel and Cunningham, 2007). While we saw significant changes between WT and HOM mice in their locomotor response to EtOH in Experiment 2, we did not see enhanced locomotor activity from the HOM mice during the five-minute conditioning sessions to EtOH or during the drug-free preference tests. Moreover, we only saw enhanced locomotor activity in the HOMs during the pretest session, consistent with a novelty-induced hyperlocomotion. Additionally, only the HOM males showed significant EtOH CPP, so it does not appear that any potential differences in baseline locomotor activity were sufficient to interfere with preference expression. Together, these findings suggest that the observed CPP was not due to EtOH-induced locomotor activation, but rather associative learning.

There are several brain regions implicated in CPP, including the NAc, HPC, BNST, and the prelimbic cortex (Pati et al., 2019b; Zheng et al., 2025; Zhou et al., 2019). *Dpp6* is highly expressed in the HPC and is one of two auxiliary β-subunits that modulates the expression and function of A-type voltage-gated Kv4.2 channels, which are important for neuronal excitability (Clark et al., 2008; Kise et al., 2021). Thus, a potential hypothesis is that increased dendritic excitability in the HPC and, therefore, increased excitatory projections to the NAc could be driving the differences in the expression of EtOH CPP (Trouche et al., 2019); however, other brain regions co-expressing *Dpp6* and Kv4.2 channels, such as the striatum or cerebellar cortex should be considered (Clark et al., 2008). Additionally, in *Dpp6* KO mice, Lin et al. (2018) found that KO mice displayed greater locomotor activity and deficits in several spatial learning and memory paradigms, such as the Morris Water Maze and Spatial Object Recognition. Similarly, Kiselycznyk, Hoffman, and Holmes (2012) found that Kv4.2 KO mice displayed greater locomotor activity and greater anxiolytic behavior, which is consistent with our results. Together, these findings support the notion that both *Dpp6* and Kv4.2 channels are co-expressed (Clark et al., 2008), and suggests that both are also important for changes in locomotor activity; however, future studies should evaluate the relationship between *Dpp6* and Kv4.2 channels to confirm their relationship and how, collectively, they might contribute to alcohol-related phenotypes.

We also found that HOMs displayed greater novelty-induced hyperactivity in the CPP chamber and the open field chamber. This behavioral response involves the HPC, VTA, and locus coeruleus among other brain regions (Fois et al., 2022; Procaccini et al., 2011). Hippocampal circuits play an important role in novelty recognition (Fanselow and Dong, 2010), which further suggests that deletion of this gene disrupts hippocampal neurotransmission that may be important for this behavior. This is in line with Procaccini et al. (2011) who found more c-Fos expression in the hippocampus of GluA1-KO mice following novelty-induced hyperactivity.

Additionally, studies of both *Dpp6* KO and Kv4.2 KO models have found that anxiety-like behavior is reduced measured by time spent in the open arms of the elevated plus maze (Kiselycznyk et al., 2012; Lin et al., 2018). We did not detect significant differences in anxiety-like behavior as measured by percent of the time spent in the center on habituation day 1 when mice received an injection of saline, which is in line with unpublished data reported by Lin et al. (2018). However, this study demonstrated that a 1.5 and 2 g/kg dose of EtOH significantly decreased anxiety-like behavior in male and female HOMs compared to other genotypes, as measured by the percent of time they spent in the center of the open field chamber. The ventral HPC (vHPC) is implicated in anxiety-related behaviors (Fanselow and Dong, 2010). In fact, lesions and chemogenetic inhibition of the vHPC have shown to decrease anxiety-like behavior (Parfitt et al., 2017; Wang et al., 2019), suggesting that the reduction in anxiety-like behavior following EtOH seen in our experiment might be due to disruption of the ventral, but not dorsal, HPC, though this is speculative. EtOH has been shown to produce anxiolytic behavior independent of locomotor increases (Boerngen-Lacerda and Souza-Formigoni, 2000), consistent with our findings in the HOM animals. This suggests that full deletion of *Dpp6* exacerbates the anxiolytic effect potentially produced by EtOH, but not necessarily due to changes in locomotor activity.

Lastly, we found greater total locomotor activity in HOMs compared to other genotypes on all test days except following a 2 g/kg EtOH dose, suggesting that this dose was sufficient to blunt their hyperlocomotor activity. It is possible that the greater center time of the HOM animals in the open field test is due at least in part to hyperactivity, though this not supported by the 2 g/kg test day data where there were no genotypic differences in activity, but the HOMs still showed greater center time compared to the other genotypes. Further, the sedative response found in the HOMs in the first 30 minutes in Experiment 2 suggests that they may be more sensitive to the sedative properties of EtOH than WT or HET mice. Importantly, this finding does not change our interpretation of the ethanol-induced anxiolysis seen in the HOMs because we saw similar findings when we focused our analysis to the first five minutes, which is expected to be when the stimulatory effects of this does are present (Bocarsly et al., 2019). While we did not see a significant stimulatory response at the 1.5 g/kg dose in any genotype, it remains to be seen if differences might emerge at a lower dose. Testing additional EtOH doses in the future would help to resolve whether the HOMs have an increased sensitivity specifically to the sedative effects of EtOH, or whether our results are due to an overall shift in their EtOH locomotor dose-response curve. Regardless, the development of significant EtOH CPP in the HOM males suggests that they are sensitive to the reinforcing properties of ethanol even in the absence of a locomotor stimulatory response at the 2 g/kg dose. Finally, the effects reported here were all seen following injected EtOH, and it will be important for future work to examine voluntary intake to determine if *Dpp6* has a role in EtOH consumption as well.

## Conclusion

*DPP6* has been implicated in several alcohol-related phenotypes in humans, which warrants further investigation. This was the first study examining the role of *Dpp6* in the rewarding and locomotor stimulatory and sedative properties of alcohol. Overall, we found that male, but not female, HOMs are more sensitive to both the rewarding properties and sedative effects of 2.0 g/kg injection dose. Anxiolytic behavior following EtOH injections of both 1.5 and 2.0 g/kg was seen in both male and female HOMs compared to HETs and WT, and both sexes show a greater novelty-induced locomotor response than the other genotypes. Future work focusing on the underlying mechanisms are necessary to understand the behavioral differences seen here and their potential relationship to risk for AUD.

## Acknowledgements

The authors wish to express sincere gratitude to Rebeka Sultana for maintaining the breeding colony for this research.

## Author Contributions

Maribel Hernandez: writing, data collection, conceptualization, methodology, and data analysis. Amanda Barkley-Levenson: writing – review and editing, conceptualization, supervision, and funding acquisition.

## Ethics approval statement

The animal studies protocols were approved by the Institutional Animal Care and Use Committee and the Animal Resource Facility at the University of New Mexico Health Sciences Center

## Data availability statement

The data that support the findings of this study are available from the corresponding author upon reasonable request.

## Permission to reproduce material from other sources

This is an open access article under the terms of the Creative Commons Attribution License, which permits use, distribution and reproduction in any medium, provided the original work is properly cited

