## Supplemental Material for "*Dpp6* Knockout Mice Exhibit Increased Ethanol Conditioned Place Preference and Acute Ethanol-Induced Anxiolytic Behavior"

### **Supplemental Analyses**

No Significant Genotype Differences in Body Weight

Weights were tracked on testing days to calculate injection volume. A three-way RM ANOVA of day, sex, and genotype revealed a significant main effect of day, [F(4.45, 266.67) = 9.36, p < .001], and sex, [F(1, 30) = 46.50, p < .001]. This indicates that there are no significant differences between genotypes, differing from prior reports in *Dpp6* KO mice (Figure S1) (8).

Locomotor Activity Across Days

Total locomotor activity across days can be seen in Figure S2. A three-way RM ANOVA of day, sex, and genotype revealed a main effect of day, [F(1.86, 52.15) = 102.31, p < .001], sex, [F(1, 28) = 13.35, p = .001], and genotype, [F(2, 28) = 11.63, p < .001]. There was also a significant interaction of day*genotype, [F(3.73, 52.15) = 5.68, p < .001]. To probe the interaction, we collapsed by sex and ran a two-way RM ANOVA of day and genotype. This analysis revealed a significant main effect of day, [F (1.939, 60.12) = 105.3, p < .001], genotype, [F (2, 31) = 7.645, p < .001], and a significant interaction of day*genotype, [F (3.879, 60.12) = 5.534, p = .002]. Next, a Tukey's multiple comparisons test indicated a significant difference between WTs and HETs (p < .05) on day 1 only, and WTs and HOMs on days 1, 2, and 3 (p's < .05). Additionally, all genotypes exhibited significant differences in locomotor activity between day 1 and day 2 and day 1 (p's < .01) and day 10 (p's < .01). HET and HOMs also exhibit significantly different locomotor activity between day 2 and day 10 (p's < .01) and day 3 and day 10 (p's < .01). However, none of the genotypes exhibited an acute stimulatory response to a 1.5 g/kg injection as measured by locomotor differences between day 2 and day 3 (p's > .05), such that day 3 activity would be significantly increased compared to day 2.

Locomotor Activity on Day 10

We did not see a significant difference across time during the initial 30 minutes following a 2 g/kg EtOH injection. To determine potential genotypic differences in recovery from the sedative effects of the 2 g/kg EtOH dose, we assessed the last 30 minutes of the session separately. A two-way ANOVA of sex and genotype revealed a significant main effect of genotype, [F(2, 32) = 4.71, p = .016]. Tukey's post hoc test indicated significant differences between female HOM and WT mice (p = .016) as well as female HOM and HET mice (p = .013). There was also a significant difference between female HOMs and male HOMs. Together, these data suggest that HOM female mice might recover from the sedative effects of EtOH more quickly than the other groups (Figure S3A).

A three-way RM ANOVA of time, sex, and genotype on the first 30 minutes compared to the last 30 minutes revealed a significant main effect of time, [F(1, 29) = 198.061, p < .001], and sex, [F(1, 29) = 8.08, p = .01]. There were also significant interactions of time*sex, [F(1, 29) = 5.25, p = .029], time*genotype, [F(1, 29) = 4.0, p = .029], and time*sex*genotype, [F(2, 29) = 4.54, p = .019]. To follow up the three-way interaction, a two-way ANOVA of sex and genotype for each 30-minute time block was used. During the first 30 minutes, there was a significant main effect of sex, [F(1, 29) = 10.74, p = .003], suggesting that females exhibit greater locomotor activity than males. During the second 30-minute time block, this analysis revealed a significant main effect of genotype, [F(2, 29) = 4.97, p = .014]. A Tukey's post hoc analysis indicated that this was driven by a significant difference between WT and HOM mice (p = .024; Figure S4B).

Lastly, a two-way ANOVA of sex and genotype on the full 60-minute session revealed a main effect of sex, [$F(1, 29) = 8.08$, $p = .01$], and no other significant interactions, suggesting that females move significantly more than males regardless of genotype (Figure S3B).

No Significant Genotype Differences in Body Weight During the Open Field Test

A three-way RM ANOVA of day, sex, and genotype revealed a significant main effect of day, [$F(2.49, 72.18) = 9.27$, $p < .001$], sex, [$F(1, 29) = 40.63$, $p < .001$], and a trend towards a main effect of genotype ($p = .06$). These data suggest HOMs, particularly males, may show mildly reduced weight compared to the other genotypes, though this difference did not reach the level of significance in either experiment (Figure S3C).

**Supplemental Figures**

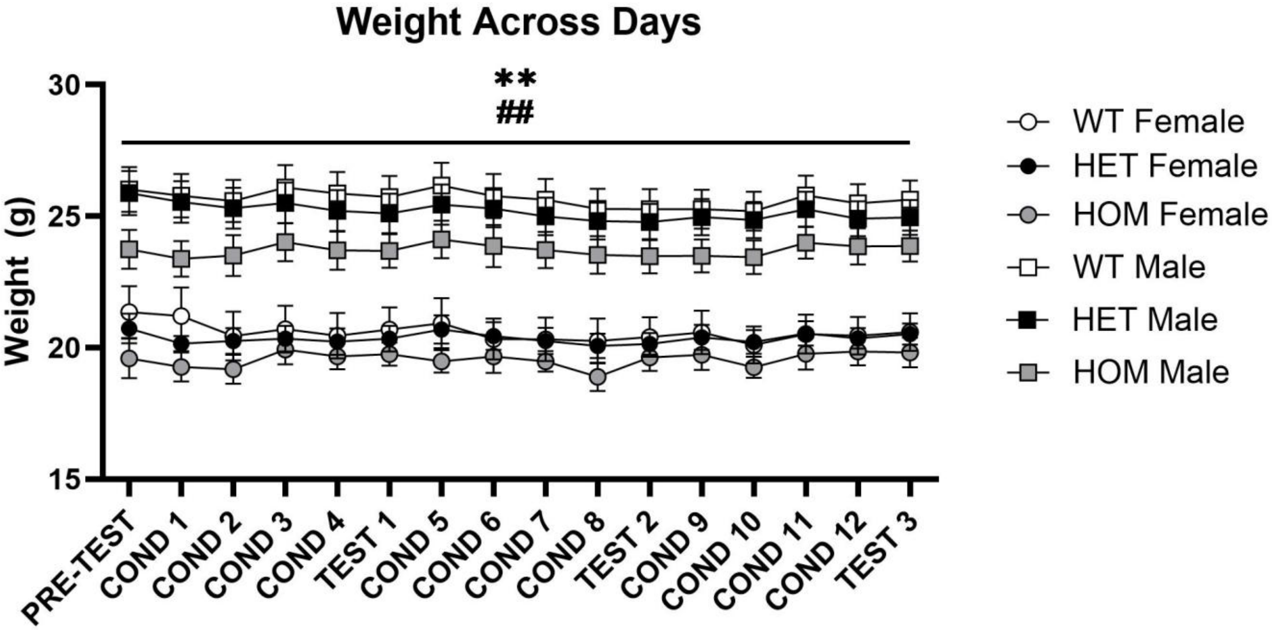


Supplemental Figure 1. Weights across injections days. There was a significant effect of day, suggesting fluctuations in weight across days. Additionally, there was a significant effect of sex, indicating that males weighed more than females. (**$p < .01$, ME day) ($^{\#\#}p < .01$, ME sex)

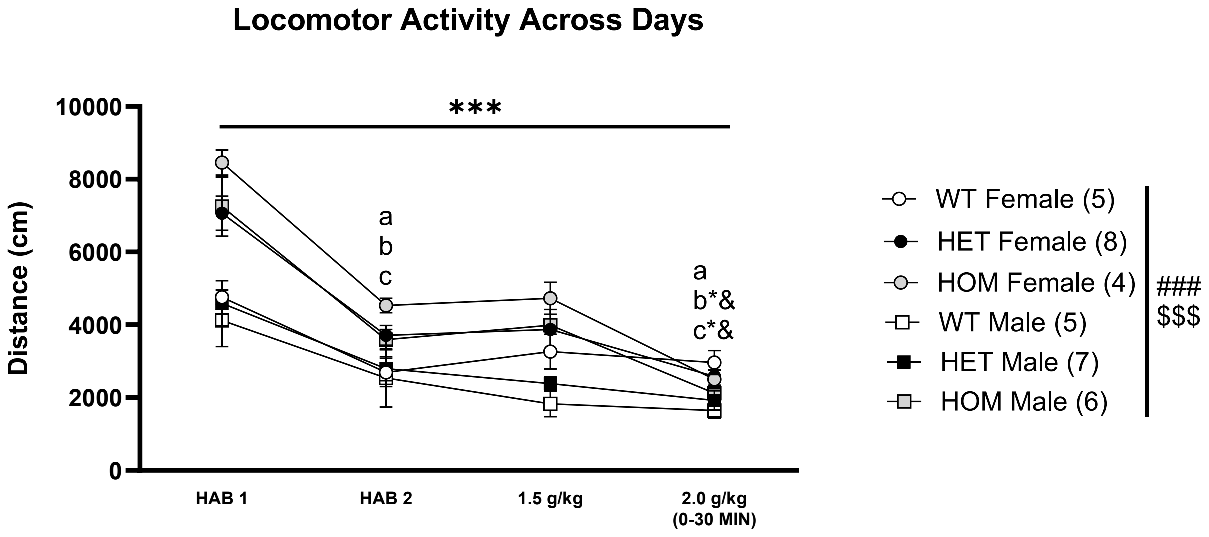


Supplemental Figure S2. Locomotor activity across days during the 30-minute session. Sex and genotype differences were detected following saline and EtOH injections. ($^{***}p < .001$ ME day; $^{\#\#\#}p < .001$ ME sex; $^{\$\$\$}p < .001$ ME Genotype; $^{a}p < .01$ WT different from day 1; $^{b}p < .01$ HET different from day 1; $^{c}p < .01$ HOM different from day 1; $^{*}p < .01$ different from day 2; $^{\&}p < .01$ different from day 3).

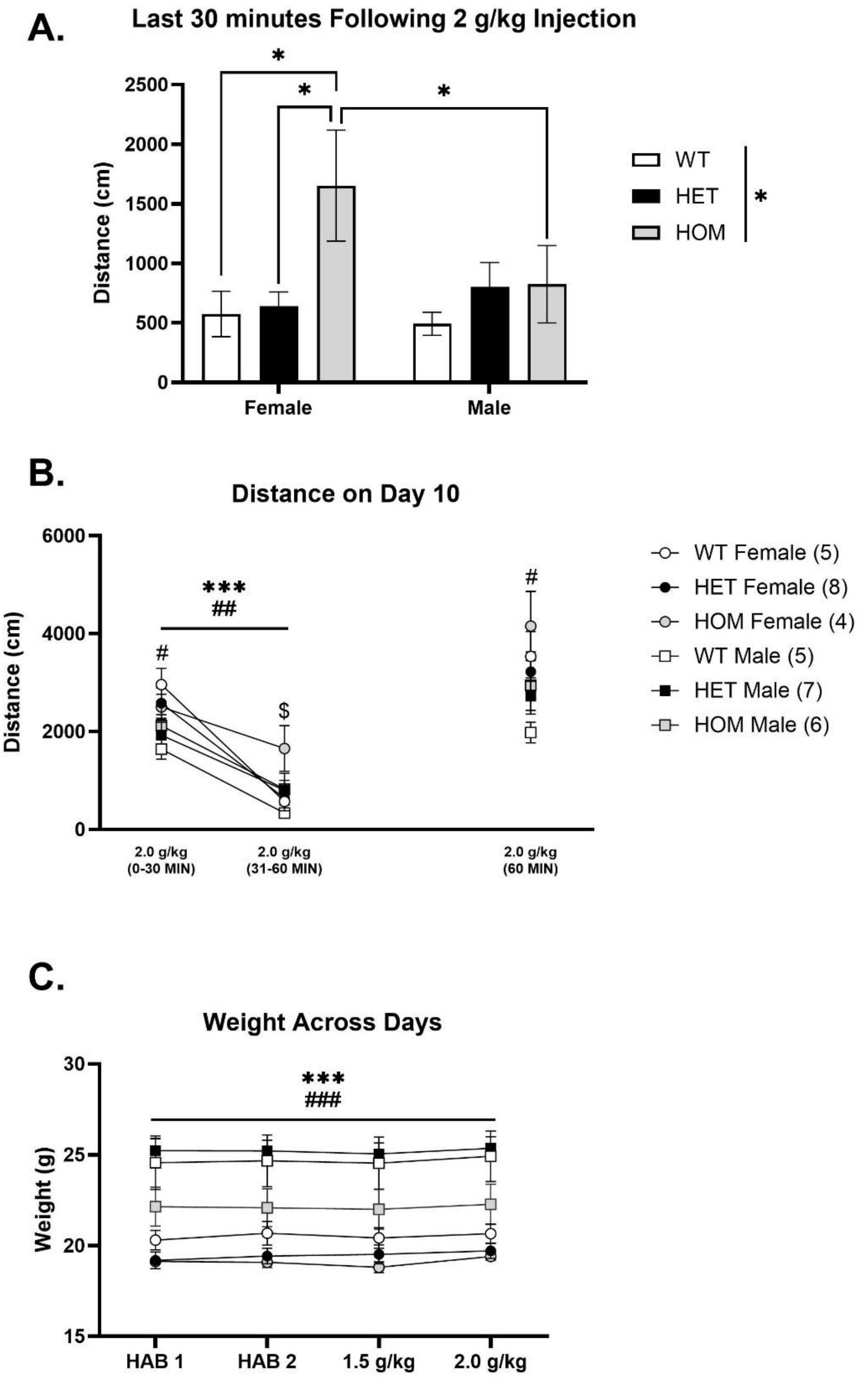


Supplemental Figure 3. Locomotor activity across the entire 60-minute session on day 10 (2 g/kg EtOH) and weights across testing days. A) Females exhibited greater locomotor activity than males across the second 30-minute time block. B) All groups decreased their locomotor activity across time blocks and females were significantly different than males during the first

30-minute time block, while this effect was not seen in the second half. HOMs exhibited greater locomotor activity compared to WTs during the second half. C). There were no significant differences in weight between different genotypes; however, males weighed more than females and there was a trend toward genotype differences ($p = .06$). (***$p < .001$ ME day; ###$p < .001$ ME Sex)
